# Reciprocal monoallelic expression of ASAR lncRNA genes controls replication timing of human chromosome 6

**DOI:** 10.1101/732784

**Authors:** Michael Heskett, Leslie G. Smith, Paul Spellman, Mathew J. Thayer

## Abstract

DNA replication occurs on mammalian chromosomes in a cell-type distinctive temporal order known as the replication timing program. We previously found that disruption of the noncanonical lncRNA genes *ASAR6* and *ASAR15* results in delayed replication timing and delayed mitotic chromosome condensation of human chromosome 6 and 15, respectively. *ASAR6* and *ASAR15* display random monoallelic expression, and display asynchronous replication between alleles that is coordinated with other random monoallelic genes on their respective chromosomes. Disruption of the expressed allele, but not the silent allele, of *ASAR6* leads to delayed replication, activation of the previously silent alleles of linked monoallelic genes, and structural instability of human chromosome 6. In this report, we describe a second lncRNA gene (*ASAR6-141*) on human chromosome 6 that when disrupted results in delayed replication timing in *cis*. *ASAR6-141* is subject to random monoallelic expression and asynchronous replication, and is expressed from the opposite chromosome 6 homolog as *ASAR6*. ASAR6-141 RNA, like ASAR6 and ASAR15 RNAs, contains a high L1 content and remains associated with the chromosome territory where it is transcribed. Three classes of *cis*-acting elements control proper chromosome function in mammals: origins of replication, centromeres; and telomeres, which are responsible for replication, segregation and stability of all chromosomes. Our work supports a fourth type of essential chromosomal element, “Inactivation/Stability Centers”, which express ASAR lncRNAs responsible for proper replication timing, monoallelic expression, and structural stability of each chromosome.

**Author summary:** Mammalian cells replicate their chromosomes during a highly ordered and cell type-specific program. Genetic studies have identified two long non-coding RNA genes, *ASAR6* and *ASAR15*, as critical regulators of the replication timing program of human chromosomes 6 and 15, respectively. There are several unusual characteristics of the ASAR6 and ASAR15 RNAs that distinguish them from other long non-coding RNAs, including: being very long (>200 kb), lacking splicing of the transcripts, lacking polyadenylation, and being retained in the nucleus on the chromosomes where they are made. *ASAR6* and *ASAR15* also have the unusual property of being expressed from only one copy of the two genes located on homologous chromosome pairs. Using these unusual characteristics shared between *ASAR6* and *ASAR15*, we have identified a second ASAR lncRNA gene located on human chromosome 6, which we have named *ASAR6-141*. *ASAR6-141* is expressed from the opposite chromosome 6 homolog as *ASAR6*, and disruption of the expressed allele results in delayed replication of chromosome 6. ASAR6-141 RNA had previously been annotated as vlinc273. The <u>v</u>ery long intergenic <u>n</u>on-<u>c</u>oding (vlinc)RNAs represent a recently annotated class of RNAs that are long (>50 kb), non-spliced, and non-polyadenlyated nuclear RNAs. There are currently >2,700 vlincRNAs expressed from every chromosome, are encoded by >15% of the human genome, and with a few exceptions have no known function. Our results suggest the intriguing possibility that the vlinc class of RNAs may be functioning to control the replication timing program of all human chromosomes.

## Introduction

Numerous reports over the past 50+ years have described an abnormal DNA replication phenotype affecting individual chromosomes in mitotic preparations from mammalian cells [1]. For example, we found that certain tumor derived chromosome translocations display a <u>d</u>elay in replication timing (DRT) that is characterized by a >3 hour delay in the initiation and completion of DNA synthesis along the entire length of individual chromosomes [2]. Chromosomes with DRT also display a <u>d</u>elay in <u>m</u>itotic chromosome <u>c</u>ondensation (DMC), which is characterized by an under-condensed appearance during mitosis and a concomitant delay in the mitotic phosphorylation of histone H3 [2, 3]. We have also found that ∼5% of chromosomal translocations induced by exposing human cells to ionizing radiation (IR) display DRT/DMC [4]. To characterize the DRT/DMC phenotype further, we developed a Cre/loxP system that allowed us to create chromosome translocations in a precise and controllable manner [4, 5]. Using this Cre/loxP system, we carried out a screen in human cells designed to identify loxP integration sites that generate chromosome translocations with DRT/DMC [4–7]. We found that ∼5% of Cre/loxP induced translocations display DRT/DMC [5]. Therefore, ∼5% of translocations induced by two different mechanisms (IR or Cre/loxP) result in DRT/DMC.

Our Cre/loxP screen identified five cell lines that generate balanced translocations, affecting eight different autosomes, all displaying DRT/DMC [5]. Characterization of two of these translocations identified discrete *cis*-acting loci that when disrupted result in DRT/DMC on human chromosomes 6 or 15 [6, 7]. Molecular examination of the disrupted loci identified two lncRNA genes, which we named <u>AS</u>ynchronous replication and <u>A</u>utosomal <u>R</u>NA on chromosome <u>6</u> (*ASAR6*) and on chromosome <u>15</u> (*ASAR15*) [6, 7]. These studies defined the first *cis*-acting loci that control replication timing, monoallelic gene expression, and structural stability of individual human autosomes [6, 7].

The vast majority of genes on mammalian autosomes are expressed from both alleles. However, some autosomal genes are expressed preferentially from only one allele, achieving a state of “autosome pair non-equivalence” [8, 9]. The most extreme form of differential allelic expression is often referred to as monoallelic expression, where a single allele is expressed exclusively (reviewed in [10]). Differential allelic expression can arise from distinct mechanisms. For example, differential expression can arise due to DNA sequence polymorphisms within promoter or enhancer elements that influence the efficiency with which a gene will be transcribed (reviewed in [11, 12]). In contrast, differential expression can occur in the absence of DNA sequence polymorphisms and is connected to situations where there is a “programmed” requirement to regulate gene dosage or to provide exquisite specificity (reviewed in [11, 13–15]). One well established form of programmed monoallelic expression occurs in a parent of origin specific manner, and is known as genomic imprinting (reviewed in [16]). In addition, monoallelic expression occurring in a random manner has been observed from as many as 8% of autosomal genes [12, 17]. One unusual characteristic of all programmed monoallelic genes is asynchronous replication between alleles [8, 9, 18, 19]. This asynchronous replication is present in tissues where the genes are not transcribed, indicating that asynchrony is not dependent on transcription [7–9, 20]. Furthermore, asynchronous replication of random monoallelic genes is coordinated with other random monoallelic genes on the same chromosome, indicating that there is a chromosome-wide system that coordinates replication asynchrony of random monoallelic genes [7–9, 20]. We use the following criteria to classify genes as being subject to <u>P</u>rogramed <u>R</u>andom <u>M</u>onoallelic <u>E</u>xpression (PRME): 1) monoallelic expression is detected in multiple unrelated individuals, which rules out rare DNA polymorphisms in promoters or enhancers; 2) monoallelic expression of either allele is detected in single cell-derived subclones from the same individual, which rules out genomic imprinting; and 3) asynchronous replication is present and coordinated with other random monoallelic genes on the same chromosome, indicating that the monoallelic gene is regulated by a chromosome-wide system that coordinates asynchronous replication along chromosome pairs. Using these criteria, we previously found that *ASAR6* and *ASAR15* are subject to PRME [6, 7, 20].

Recent reports have described <u>v</u>ery long intergenic <u>n</u>on-<u>c</u>oding (vlinc)RNAs expressed in numerous human tissues [21–23]. The vlincRNAs are RNA Pol II products that are nuclear, non-spliced, non-polyadenylated transcripts of >50 kb of contiguously expressed sequence that are not associated with protein coding genes. The initial reports annotated 2,147 human vlincRNAs from 833 samples in the FANTOM5 dataset [23, 24]. A more recent study identified an additional 574 vlincRNAs expressed in childhood acute lymphoblastic leukemia [25]. Therefore, there are currently >2,700 annotated vlincRNAs that are encoded by >15% of the human genome [23–25]. ASAR6 and ASAR15 RNAs share several characteristics with the vlincRNAs, including: RNA Pol II products, long contiguous transcripts (>50 kb) that are non-spliced, non-polyadenylated, and are retained in the nucleus [6, 7, 20]. Therefore, given these shared characteristics between ASAR6, ASAR15 and vlincRNAs, we consider the vlincRNAs as potential ASAR candidates.

ASAR6 and ASAR15 RNAs also share additional characteristics, including: PRME, are retained within the chromosome territories where they are transcribed, and contain a high long interspersed element 1 (LINE1 or L1) content [6, 7, 20]. In this report, we used these “ASAR” characteristics to identify a second lncRNA gene that controls replication timing of human chromosome 6, which we designate as *ASAR6-141*. The *ASAR6-141* gene is located at ∼141 mb of human chromosome 6, is subject to random monoallelic expression and asynchronous replication, and disruption of the expressed allele, but not the silent allele, leads to delayed replication of human chromosome 6 in *cis*. ASAR6-141 RNA, which was previously annotated as vlinc273 [23], is ∼185 kb in length, contains ∼30% L1 sequences, remains associated with the chromosome 6 territory where it is transcribed, and is expressed in *trans* to the expressed allele of *ASAR6*. These observations support a model that includes reciprocal monoallelic expression of different ASAR lincRNA genes that control replication timing of homologous chromosome pairs [26].

## Results

### Reciprocal random monoallelic expression of ASAR lncRNAs on chromosome 6

With the goal of identifying nuclear RNAs with “ASAR” characteristics expressed from human chromosome 6, we carried out RNA-seq on nuclear RNA isolated from HTD114 cells. HTD114 cells are a human fibrosarcoma cell line, where we previously carried out the Cre/loxP screen that led to the identification and functional characterization of *ASAR6* and *ASAR15* [5–7]. Figure 1a shows the UCSC Genome Browser view of chromosome 6, between 140.2 mb and 141.3 mb, showing the RNA-seq reads from the region previously annotated as expressing 6 different vlincRNAs [23]. We note that vlinc273 is expressed, but vlinc271, vlinc1010, vlinc1011, vlinc1012, and vlinc272, show little or no expression in HTD114 cells. Also shown in Figure 1 are the location of Fosmids used in the RNA-DNA FISH analyses (see below), the Long RNA-seq track (showing contiguous transcripts from three human cell lines, GM12878, HepG2, and K562) from ENCODE/Cold Spring Harbor, and the Repeat Masker Track showing the location of repetitive elements (also see Table S1).

**Fig 1.**
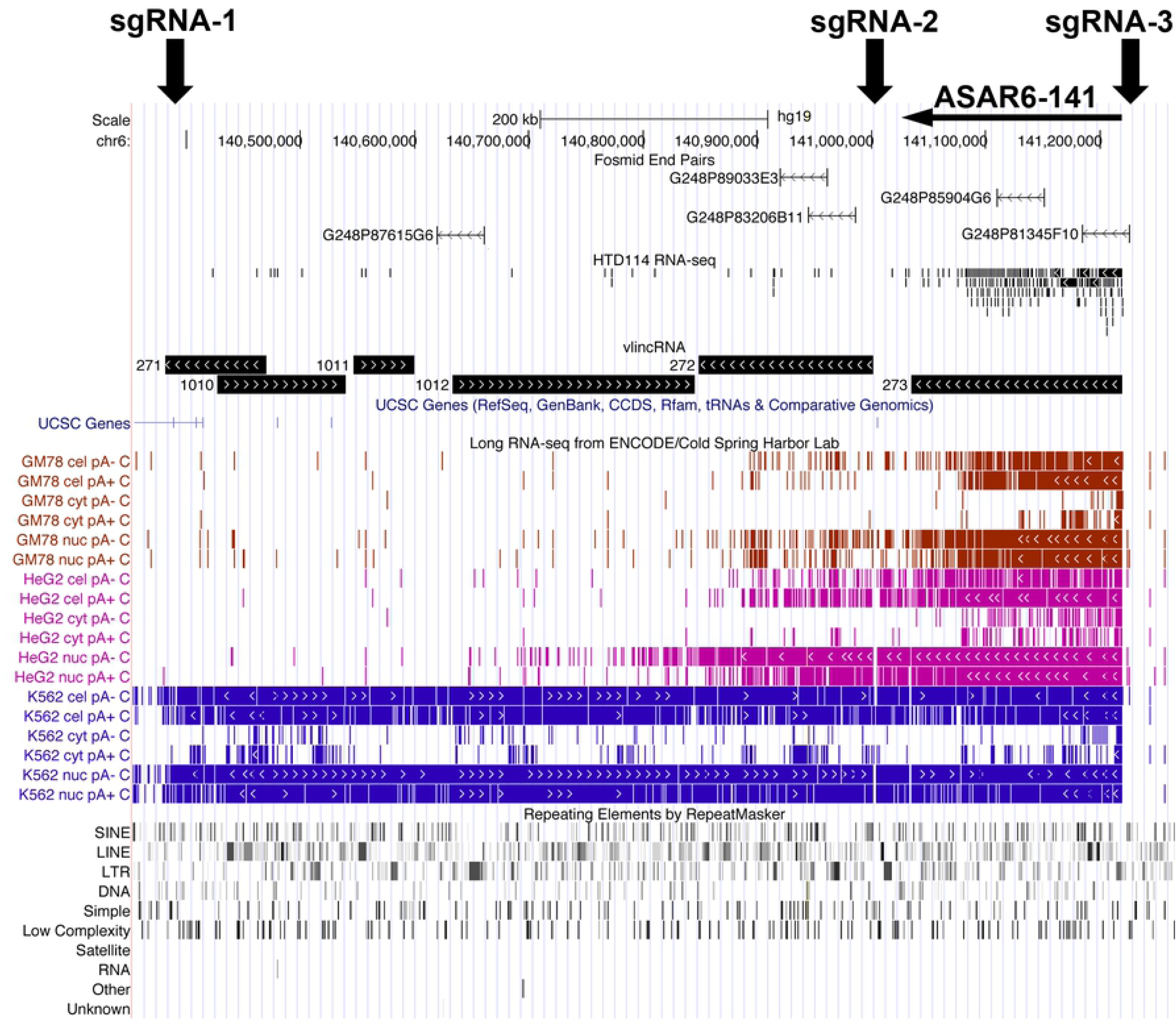
UCSC Genome Browser view of the vlinc cluster on chromosome 6 between 140.3 and 141.3 mb. The genomic locations of vlinc271, vlinc1010, vlinc1011, vlinc1012, vlinc272 and vlinc273 are illustrated using the UCSC Genome Browser. RNA-seq data from nuclear RNA isolated from HTD114 is shown (HTD114 RNA-seq). Long RNA-seq data from the ENCODE Project (Cold Spring Harbor Lab) is shown using the Contigs view. Expression from the human cell lines GM12878 (red), K562 (dark blue), Hela S3 (light blue), and HepG2 (magenta) are shown. RNA from total cellular Poly A+ (cel pA+), total cellular Poly A-(cel pA-), nuclear Poly A+ (nuc pA+), nuclear Poly A-(nuc pA-), cytoplasmic Poly A+ (cyt pA+), and cytoplasmic Poly A-(cyt pA-) are shown. Also shown are the Repeating Elements using the RepeatMasker track. The location of five RNA FISH probes (Fosmids) that were used to detect expression of vlinc1012 (G248P87615G6), vlinc272 (G248P89033E3 and G248P83206B11) and vlinc273 (G248P85904G6 and G248P81345F10) are shown.

Next, to determine if the vlinc273 transcripts show monoallelic expression in HTD114 cells, we used reverse transcribed RNA as input for PCR, followed by sequencing at heterozygous SNPs. Figure 2a shows sequencing traces from two different SNPs that are heterozygous in genomic DNA, but a single allele was detected in RNA isolated from HTD114 cells, indicating that these transcripts are monoallelic. In addition, we previously generated two chromosome 6 mono-chromosomal hybrids to aid in mapping heterozygous SNPs onto the HTD114 chromosome 6 homologs [6]. These two hybrid cell lines are mouse L cell clones, each containing one of the two chromosome 6s from HTD114, which we arbitrarily name as CHR6A and CHR6B. Using these mono-chromosomal hybrids, we previously found that *ASAR6* is expressed from CHR6A [6]. Sequence traces generated from genomic DNA isolated from these mono-chromosomal hybrids indicated that the vlinc273 transcripts are derived from CHR6B (Fig. 2A), and therefore are expressed from the opposite chromosome, or in *trans*, to *ASAR6*.

**Fig 2.**
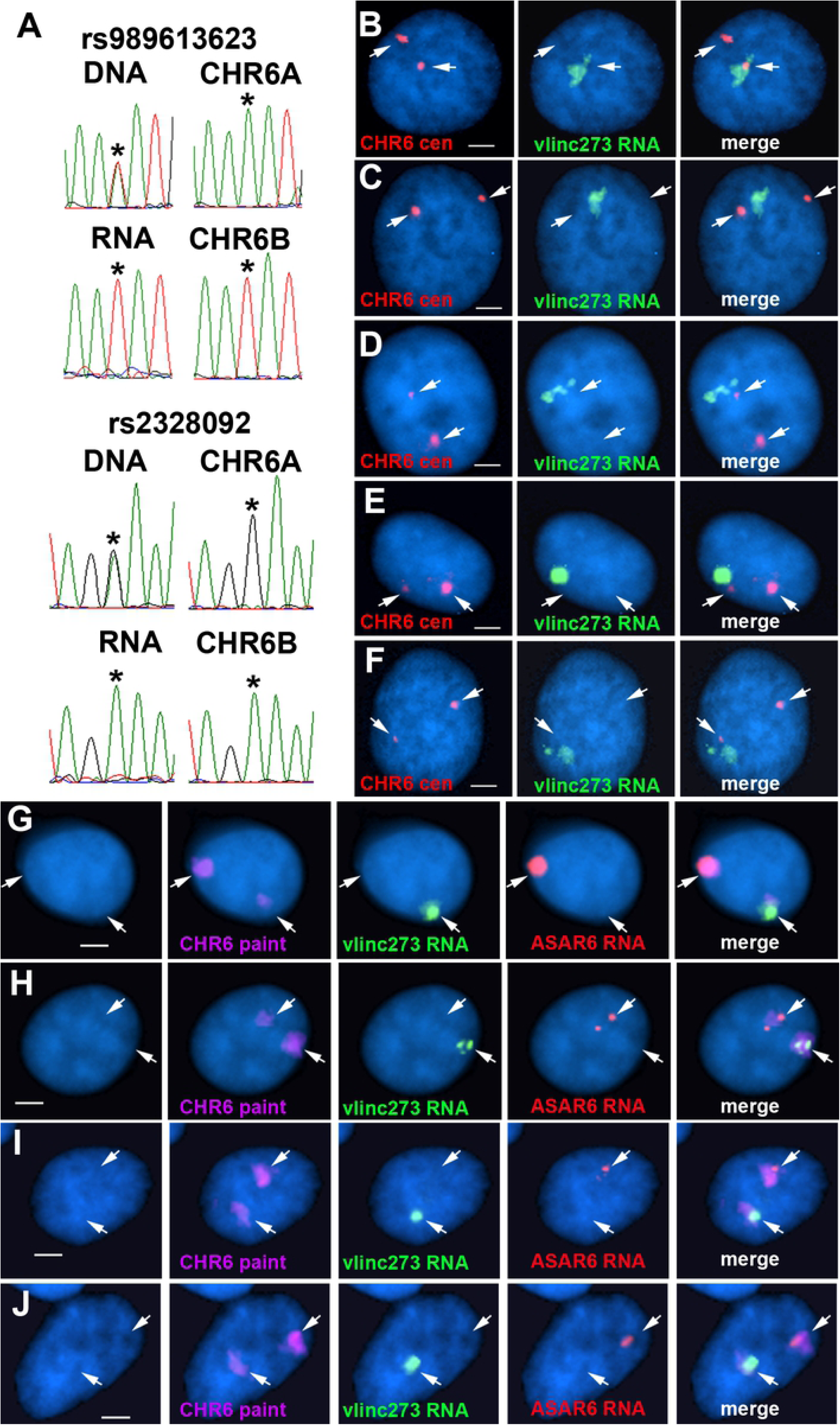
Mono-allelic expression and nuclear retention of vlinc273 in HTD114 cells. (A) DNA sequencing traces from PCR products designed to detect SNPs rs989613623 and rs2328092. PCRs were carried out on genomic DNAs isolated from HTD114, two mono-chromosomal hybrids containing the two different chromosome 6s from HTD114 {L(Hyg)-1 contains chromosome 6A (CHR6A) and expresses *ASAR6*, and L(Neo)-38 contains chromosome 6B (CHR6B) and is silent for *ASAR6* [6]}. The top and bottom panels also show the traces from HTD114 cDNA (RNA). The asterisks mark the location of the heterozygous SNPs. (B-F) RNA-DNA FISH to detect vlinc273 expression in HTD114 cells. Fosmid G248P81345F10 was used as probe to detect vlinc273 RNA (green), and a chromosome 6 centromeric probe was used to detect chromosome 6 DNA (red). The nuclear DNA was stained with DAPI. Bars are 2.5 uM. (G-J) RNA-DNA FISH to detect vlinc273 and ASAR6 expression in HTD114 cells. Fosmid G248P81345F10 was used as probe to detect vlinc273 RNA (green), Fosmid G248P86031A6 was used as probe to detect ASAR6 RNA (red), and a chromosome 6 paint was used to detect chromosome 6 DNA (magenta). The nuclear DNA was stained with DAPI. Bars are 2.5 uM.

We next assayed expression of the vlincRNA cluster using RNA-DNA FISH in HTD114 cells. For this analysis we used five different Fosmid probes to detect RNA (see Fig. 1), plus a chromosome 6 centromeric probe to detect DNA. As expected from the RNA-seq analysis, we did not detect expression from the genomic regions annotated as vlinc1012 and vlinc272 in HTD114 cells (not shown). In contrast, we detected expression of RNA, annotated as vinc273, that remains associated with one of the chromosome 6 homologs. Figure 2b-f show examples of this analysis using probes from within the vlinc273 locus to detect RNA. Note the relatively large clouds of RNA that are adjacent to, or overlapping with, one of the chromosome 6 centromeric DNA signals. In addition, we used RNA-DNA FISH to detect both ASAR6 and vlinc273 RNA in combination with a chromosome 6 whole chromosome paint as probe to detect chromosome 6 DNA. Figure 2g-j shows examples of this analysis and indicates that ASAR6 and vlinc273 RNAs are detected on opposite chromosome 6 homologs. We also note that the size of the RNA FISH signals detected by the ASAR6 and vlinc273 probes were variable, ranging from large clouds occupying the entire chromosome 6 territory, to relatively small spots of hybridization.

### Random monoallelic expression of vlinc273

The observations described above indicate that vlinc273 and ASAR6 RNAs are detected on opposite chromosome 6 homologs in the clonal cell line HTD114. This monoallelic expression could be due to either genomic imprinting, to DNA sequence polymorphisms within promoter or enhancer elements, or to PRME (see above). Therefore, to distinguish between these possibilities, we first determined if vlinc273 is monoallelically expressed in EBV transformed lymphoblasts, which have been used extensively in the analysis of autosomal monoallelic expression in humans [6, 8, 17, 20]. For this analysis, we used RNA-DNA FISH to assay expression of vlinc273 in GM12878 cells. For this analysis we used Fosmid probes to detect RNA (see Fig. 1), plus a chromosome 6 centromeric probe to detect DNA. We detected single sites of vlinc273 RNA hybridization in >95% of GM12878 cells (see Fig. 3A-E for examples).

**Fig 3.**
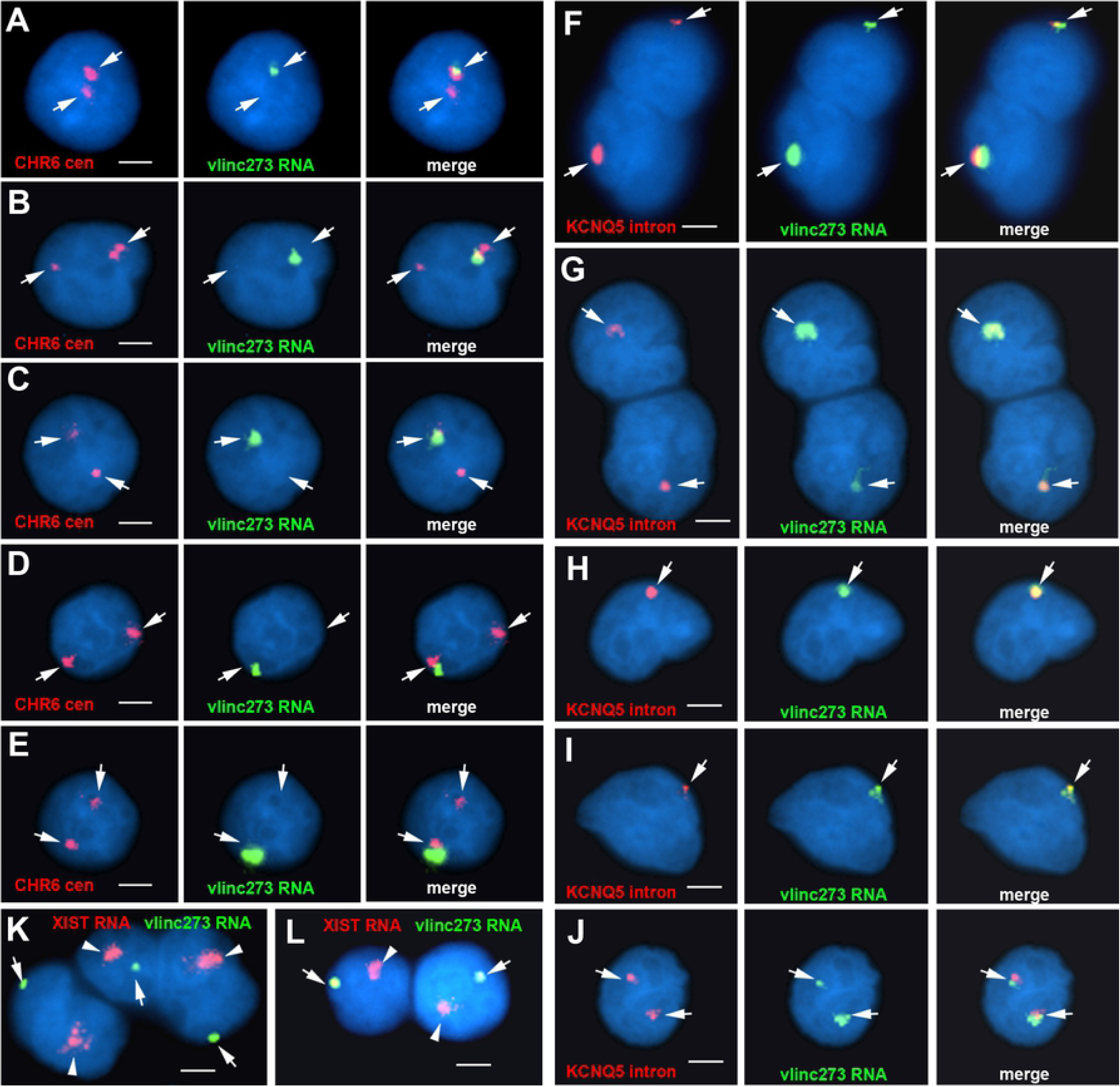
Mono-allelic expression and nuclear retention of vlinc273 in EBV transformed lymphoblasts and primary blood lymphocytes. (A-E) RNA-DNA FISH to detect vlinc273 expression in GM12878 EBV transformed lymphocytes. Fosmid G248P81345F10 was used to detect vlinc273 RNA (green), and a chromosome 6 centromeric probe (CHR6 cen) was used to detect chromosome 6 DNA (red). (F-J) RNA FISH to detect coordinated expression of vlinc273 and KCNQ5, a known random monoallelic gene, in primary blood lymphocytes. Fosmid G248P81345F10 was used to detect vlinc273 RNA (green), and Fosmid G248P80791F6 was used to detect expression of the first intron of KCNQ5. (K and I) RNA FISH to detect expression of vlinc273 and XIST RNAs, in female primary blood lymphocytes. The nuclear DNA was stained with DAPI. Bars are 2.5 uM.

Next, to determine if expression of vlinc273 is subject to PRME in human primary cells, we carried out RNA FISH on primary blood lymphocytes (PBLs) isolated from two unrelated individuals. For this analysis, we included an RNA FISH probe to the first intron of *KCNQ5*, which is a known PRME gene located on human chromosome 6 [6, 17, 20]. For this analysis, we used a two-color RNA FISH assay to detect expression of vlinc273 in combination with a probe from the first intron of *KCNQ5* on PBLs isolated from two unrelated individuals. Quantification of the number of RNA FISH signals in >100 cells indicated that vlinc273 and *KCNQ5* were expressed from the same chromosome 6 homolog in ∼98% of cells from both individuals (see Fig. 3F-I for examples). Therefore, because *KCNQ5* expression is subject to PRME [6, 17, 20], and the PBLs are not clonally derived, we conclude that the monoallelic expression of vlinc273 must also be random and therefore not imprinted. We note that the size of the RNA hybridization signals detected by the *KCNQ5* and vlinc273 probes were variable, ranging from large clouds to relatively small sites of hybridization. We also detected two sites of hybridization for both probes in ∼2% of cells (Fig. 3J). Finally, to directly compare the appearance of the RNA FISH signals detected for vlinc273 to XIST RNA expressed from the inactive X chromosome we assayed vlinc273 and XIST RNAs simultaneously in female PBLs. Figure 3k and 3i show the clouds of RNA detected by the vlinc273 probe in relation to the relatively larger clouds of RNA hybridization detected by the *XIST* probe.

### Asynchronous replication of vlinc273 is coordinated on chromosome 6

All monoallelically expressed genes share the property of asynchronous replication [27]. We previously used <u>Re</u>plication <u>Ti</u>ming-<u>S</u>pecific <u>H</u>ybridization (ReTiSH) [19] to assay coordinated asynchronous replication of chromosome 6 loci, including *ASAR6*. In the ReTiSH assay, cells are labeled with BrdU for different times and then harvested during mitosis (see Fig. 4A). Regions of chromosomes that incorporate BrdU are visualized by a modification of <u>c</u>hromosome <u>o</u>rientation-fluorescence in <u>s</u>itu <u>h</u>ybridization (CO-FISH), where the replicated regions (BrdU-labeled) are converted to single stranded DNA and then hybridized directly with specific probes [19]. Since mitotic chromosomes are analyzed for hybridization signals located on the same chromosome in metaphase spreads, the physical distance between the loci is not a limitation of the ReTiSH assay [19]. We previously used this approach to show that the asynchronous replication of *ASAR6* was coordinated in *cis* or in *trans* with other random monoallelic loci on human chromosome 6 [6, 20]. For this analysis, we used PBLs and a three-color hybridization scheme to simultaneously detect the vlinc273 locus, *ASAR6*, and the chromosome 6 centromere. The chromosome 6 centromeric probe was included to unambiguously identify both chromosome 6s. Because centromeric heterochromatin is late replicating, centromeric probes hybridize to both copies of chromosome 6 at the 14 and 5 hour time points [19]. We found that the vlinc273 alleles were subject to asynchronous replication that is coordinated in *cis* with *ASAR6* (Fig. 4B-D; and Table 1). Therefore, because the asynchronous replication of *ASAR6* is coordinated with other random monoallelic loci on chromosome 6 [6, 20], we conclude that the vlinc273 locus is part of a chromosome-wide system that coordinates the asynchronous replication of random monoallelic loci on chromosome 6.

**Fig 4.**
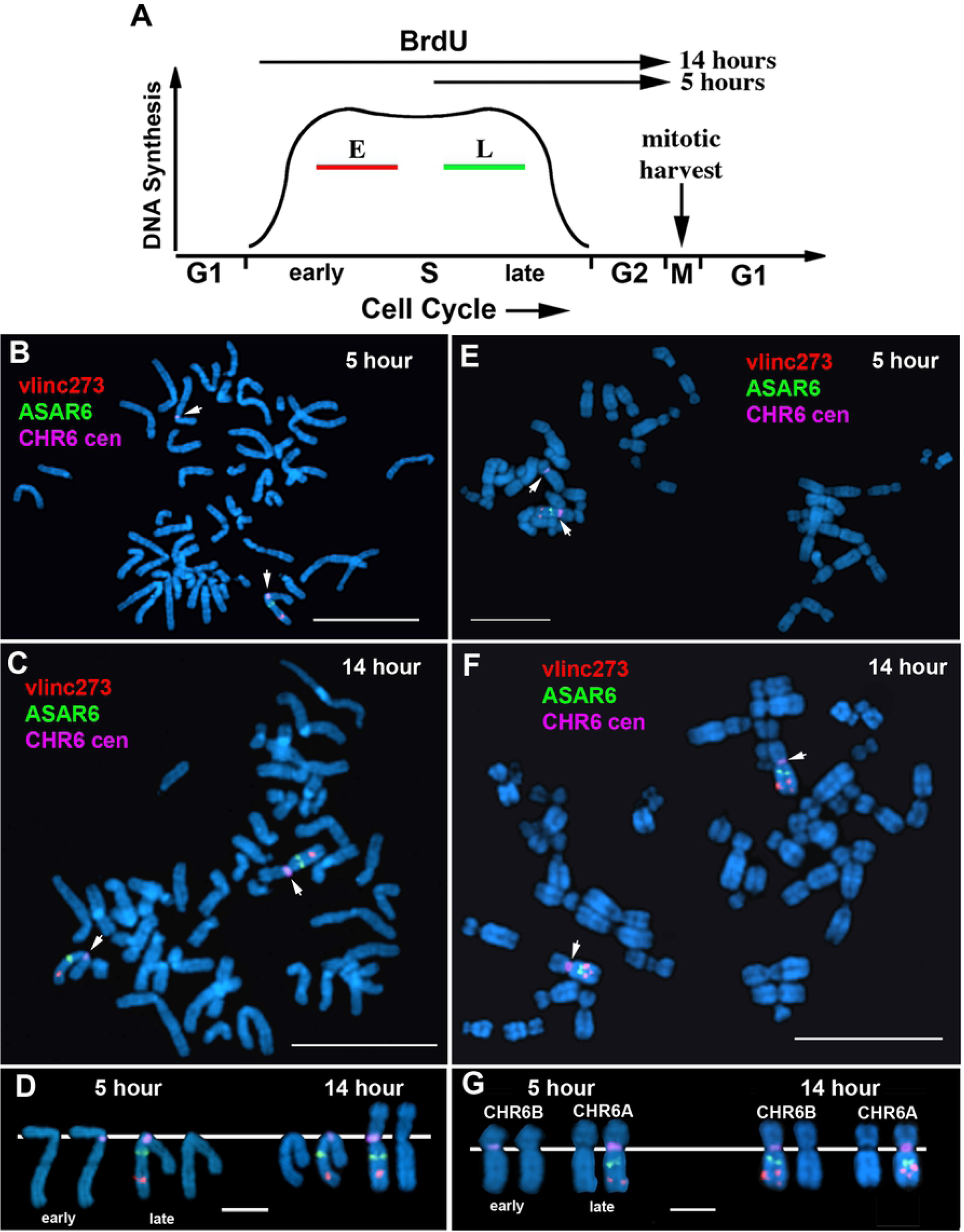
Coordinated asynchronous replication timing on chromosome 6. (A) Schematic representation of the ReTiSH assay. Cells were exposed to BrdU during the entire length of S phase (14 hours) or only during late S phase (5 hours). The ReTiSH assay can distinguish between alleles that replicate early (E) and late (L) in S phase. (B-D) Mitotic spreads that were processed for ReTiSH were hybridized with three different FISH probes. First, each hybridization included a centromeric probe to chromosome 6 (magenta). Arrows mark the centromeric signals in panels B (5 hours) and C (14 hours). Each assay also included BAC probes representing *ASAR6* (RP11-374I15; green) and vlinc273 (RP11-715D3; red). Panel D shows the two chromosome 6s, from both the 5 hour and 14 hour time points, aligned at their centromeres. The *ASAR6* BAC and the vlinc273 BAC show hybridization signals on the same chromosome 6 at the 5-hour time point, and as expected hybridized to both chromosome 6s at the 14 hour time point. The chromosomal DNA was stained with DAPI. (E-G) ReTiSH assay on HTD114 cells. Each ReTiSH assay included a centromeric probe to chromosome 6 (magenta). Arrows mark the centromeric signals in panels e (5 hours) and f (14 hours). Each assay also included BAC probes for *ASAR6* (RP11-374I15; green) and *vlinc273* (RP11-715D3; red). The chromosomal DNA was stained with DAPI. The *ASAR6* BAC and the vlinc273 BAC show hybridization signals to the same chromosome 6s (CHR6A) at the 5-hour time point, and as expected hybridized to both chromosome 6s at the 14-hour time point.

**Table 1.** Coordinated Asynchronous Replication Timing by ReTiSH.

| **PBLs** |  |  |  |  |  |
| --- | --- | --- | --- | --- | --- |
| **Locus 1** | **Locus 2** |  | **cis (%)** | **trans (%)** | **P value** |
| ASAR6-141 | ASAR6 |  | 75 | 25 | <1 x 10<sup>-3</sup> |

| **HTD114** |  |  |  |  |  |
| --- | --- | --- | --- | --- | --- |
| **Locus 1** | **Locus 2** |  | **cis (%)** | **trans (%)** | **P value** |
| ASAR6-141 | CHR6 cen* |  | 77 | 23 | <1 x 10<sup>-4</sup> |
| ASAR6 | CHR6 cen* |  | 88 | 12 | <1 x 10<sup>-6</sup> |
| ASAR6-141 | ASAR6 |  | 76 | 24 | <1 x 10<sup>-4</sup> |

CHR6 cen\* indicates the large centromere on one of the chromosome 6s in HTD114 cells.

One shared characteristic of the *ASAR6* and *ASAR15* genes is that the silent alleles replicate before the expressed alleles on their respective chromosomes [6, 7, 20]. Therefore, one unanticipated result from our ReTiSH assay is that asynchronous replication of vlinc273 and *ASAR6* is coordinated in *cis*. Thus, the earlier replicating vlinc273 allele is on the same homolog as the earlier replicating *ASAR6* allele. Therefore, to determine if the asynchronous replication of vlinc273 and *ASAR6* is also coordinated in *cis* in HTD114 cells, where they are expressed from opposite homologs (see Fig. 2), we analyzed the asynchronous replication of vlinc273 and *ASAR6* using the same three color ReTiSH assay describe above [19]. In addition, HTD114 cells contain a centromeric polymorphism on chromosome 6, and the chromosome with the larger centromere is linked to the later replicating and expressed allele of *ASAR6* [20]. We found that the asynchronous replication of vlinc273 and *ASAR6* is coordinated in *cis* in HTD114 cells (Fig. 4E-G; and Table 1). These observations are consistent with our previous finding that *ASAR6* is expressed from the later replicating allele ([20]; CHR6A), and indicate that vlinc273 is expressed from the earlier replicating allele in HTD114 cells (CHR6B; see Fig. 2A). Regardless, we found that the vlinc273 locus is subject to random monoallelic expression and asynchronous replication that is coordinated with other random monoallelic loci on chromosome 6 and therefore vlinc273 is subject to PRME.

### Deletion of the expressed allele of vlinc273 results in delayed replication in *cis*

To determine if the genomic region containing the vlincRNA cluster located on chromosome 6 at 140.2-141.3 mb (see Fig. 1) regulates replication timing, we used CRISPR/Cas9 to delete the entire locus in HTD114 cells. For this analysis we designed single guide RNAs (sgRNAs) to unique sequences as shown in Fig 1. We expressed sgRNA-1 and sgRNA-3 in combination with Cas9 and screened clones for deletions using PCR primers that flank the sgRNA binding sites (see Fig. 1 and Table S2). Because vlinc273 expression is monoallelic in HTD114 cells (see Fig. 2), we isolated clones that had heterozygous deletions affecting either CHR6A or CHR6B. We determined which allele was deleted based on retention of the different base pairs of heterozygous SNPs located within the deleted regions (see Table S2).

From our previous studies, we knew that prior to any genetic alterations the chromosome 6 homologs replicate synchronously in HTD114 cells [5, 6, 20, 26]. In addition, we also took advantage of the centromeric polymorphism in HTD114 cells to unambiguously distinguish between the two chromosome 6 homologs ([20, 26]; see Fig. 4E-G). The chromosome 6 with the larger centromere is linked to the expressed allele of *ASAR6* ([20]; CHR6A), and therefore the expressed allele of vlinc273 is linked to the chromosome 6 with the smaller centromere (CHR6B). For this replication timing assay, cultures were incubated with BrdU for 5.5 hours and mitotic cells harvested, processed for BrdU incorporation and subjected to FISH using a chromosome 6 centromeric probe. As expected, prior to disruption of the vlinc cluster, CHR6A and CHR6B display synchronous replication (see Fig. 5F below). In contrast, cells containing a deletion of the vlinc cluster on CHR6B contain significantly more BrdU incorporation into CHR6B than in CHR6A (Fig. 5A-E). Quantification of the BrdU incorporation in multiple cells indicated that deletion of the CHR6B allele, which contains the expressed allele of vlinc273, results in a significant delay in replication timing (Fig. 5F). This is in contrast to cells containing a deletion of the vlinc cluster from the CHR6A allele, which is silent for all 6 vlincRNAs, where the BrdU incorporation is comparable between CHR6A and CHR6B (Fig. 5F). In addition, replication timing analysis of heterozygous deletions encompassing only the vlinc273 locus (using sgRNA-2 and sgRNA-3) indicated that deletion of the expressed allele (CHR6B), but not the silent allele (CHR6A), resulted in delayed replication of chromosome 6 (Fig. 5F). Finally, deletion of the vlinc271, vlinc1010, vlinc1011, vlinc1012 and vlinc272 loci (using sgRNA-1 and sgRNA-2) on CHR6B, did not result in delayed replication of chromosome 6 (Fig. 5F). For an additional comparison, we included the chromosome 6 replication timing data from HTD114 cells containing heterozygous deletions of *ASAR6* on the expressed allele (CHR6A) and on the silent allele (CHR6B) (Fig. 5F; also see Fig. S1). Taken together these results indicate that deletion of the expressed allele of vlinc273 results in delayed replication of chromosome 6 in *cis*, and because vlinc273 also displays PRME, the vlinc273 locus is an ASAR. Because vlinc273 is the second ASAR identified on human chromosome 6 and is located at ∼141 mb, we designate this gene as *ASAR6-141*.

**Fig 5.**
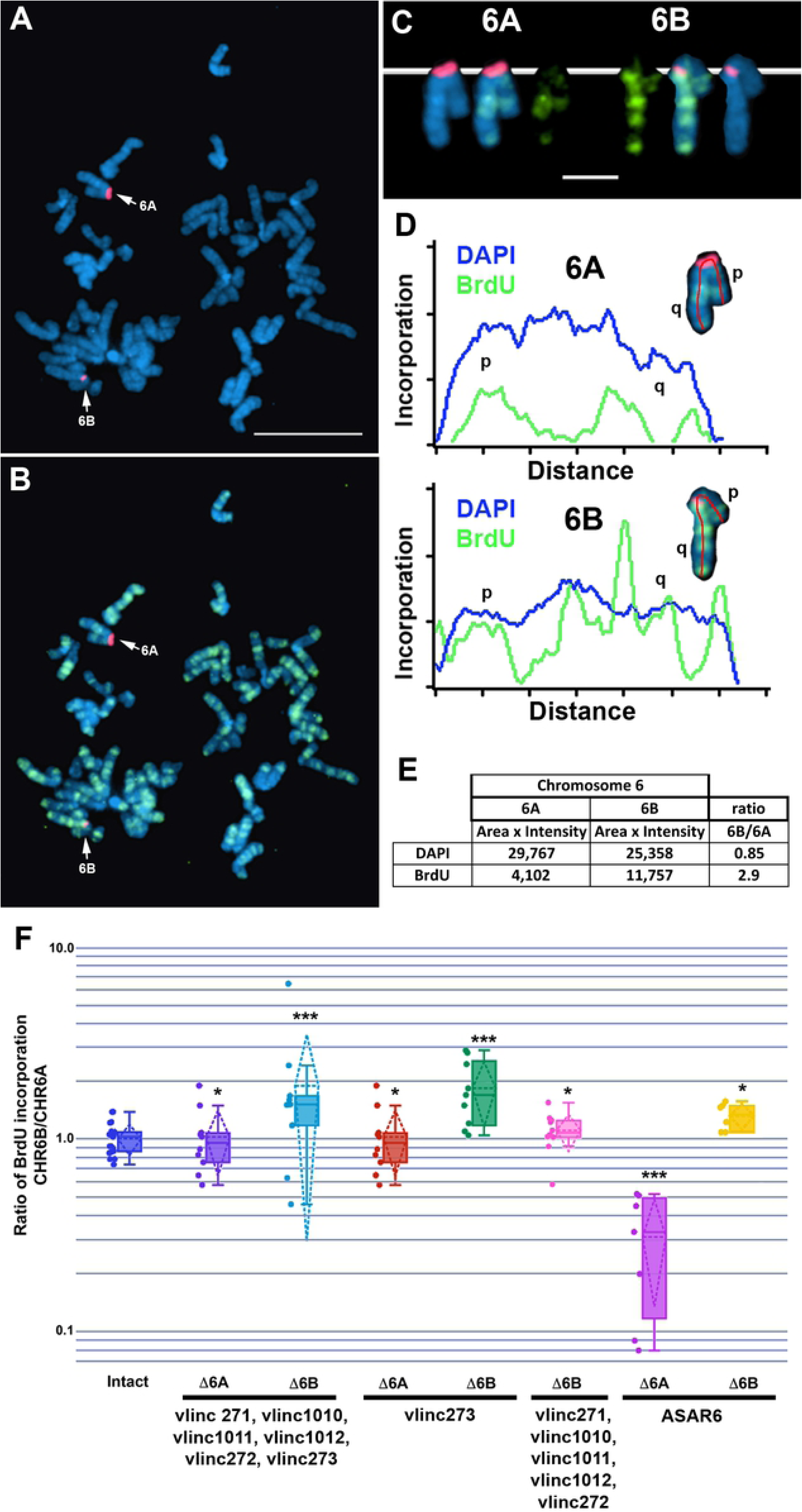
Delayed replication of chromosome 6 following disruption of vlinc273. (A and B) A representative mitotic spread from BrdU (green) treated cells containing a deletion of the expressed allele of the vlinc273 locus. Mitotic cells were subjected to DNA FISH using a chromosome 6 centromeric probe (red). The larger centromere resides on the chromosome 6 with the expressed *ASAR6* allele and the silent vlinc273 allele (6A). (C) The two chromosome 6s were extracted from a and b and aligned to show the BrdU incorporation and centromeric signals. (D) Pixel intensity profiles of BrdU incorporation and DAPI staining along the (6A) and (6B) chromosomes from panel C. (E) BrdU quantification along 6A and 6B from panel D. (F) The ratio of DNA synthesis into the two chromosome 6s was calculated by dividing the BrdU incorporation in 6B by the incorporation in 6A in multiple cells. The box plots show the ratio of incorporation before (Intact, dark blue), and heterozygous deletions of the entire locus (Δ6A purple; and Δ6B light blue), which included vlinc271, vlinc1010, vlinc1011, vlinc1012, vlinc272, and vlinc273, see map in Fig 1. Heterozygous deletions affecting vlinc273 only from the silent (Δ6A orange) or expressed (Δ6B green) alleles are shown. A heterozygous deletion affecting vlinc271, vlinc1010, vlinc1011, vlinc1012, and vlinc272 on CHR6B (Δ6B) is shown in pink. Also shown are the heterozygous deletions affecting *ASAR6* from on the expressed (Δ6A magenta) or silent (Δ6B yellow) alleles. P values of <1 x10^-4^ are indicated by ***, and P values of >1 x 10^-1^ are indicated by *, and were calculated using the Kruskal-Wallis test. Error bars are SD.

## Discussion

Chromosome associated lncRNAs have become well established as regulators of chromosome scale replication timing, gene expression and structural stability [1, 28]. In this report, we identified a second chromosome 6 lncRNA gene, *ASAR6-141*, that when disrupted results in delayed replication timing of the entire chromosome in *cis*. *ASAR6* and *ASAR6-141* are subject to PRME, are expressed from opposite chromosome 6 homologs, and disruption of the expressed alleles, but not the silent alleles, leads to delayed replication timing of human chromosome 6 in *cis*. ASAR6 and ASAR6-141 RNAs share certain characteristics, including RNA Pol II products that are non-spliced, non-polyadenylated, contain a high L1 content and remain associated with the chromosome territories where they are transcribed. Taken together our results indicate that the replication timing of human chromosome 6 is regulated by the reciprocal monoallelic expression of two different ASAR lncRNA genes (see Fig. 6).

**Fig 6.**
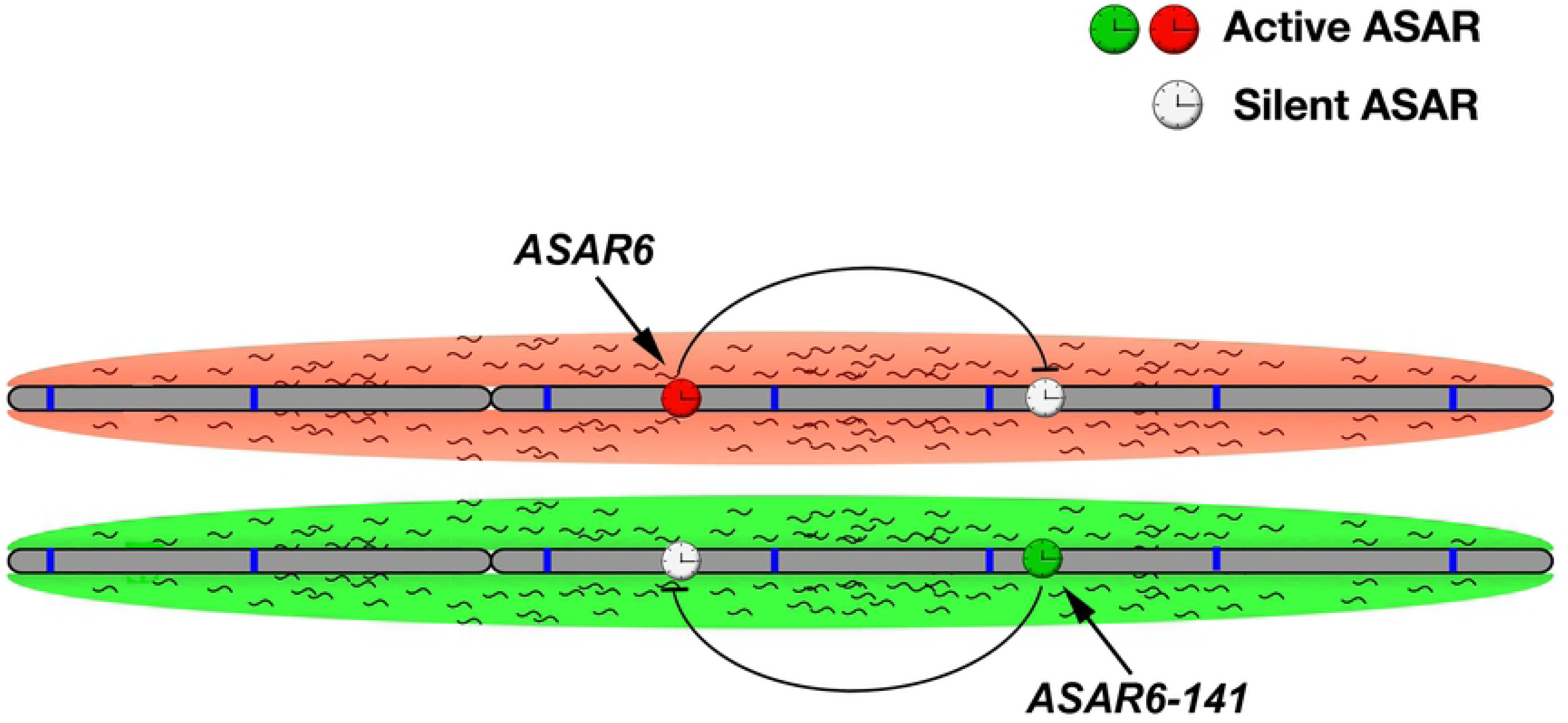
“ASAR” model of replication timing on chromosome 6. The two homologs of human chromosome 6 are shown (gray) with origins of replication depicted as blue bars. Expression of *ASAR6* and *ASAR6-141* genes is monoallelic, resulting in a reciprocal expression pattern with an expressed or active ASAR (green or red clock) and a silent or inactive ASAR (white clock) on each homolog. The red and green clouds surrounding the chromosomes represent “ASAR” RNA expressed from the different active “ASARs” on each homolog.

We previously found that deletion of the expressed allele of *ASAR6* results in transcriptional activation of the previously silent alleles of other monoallelic genes nearby, indicating that *ASAR6* negatively regulates expression of the previously silent alleles of other linked monoallelic genes [6]. One important tool for the analysis of chromosome scale gene expression has been the use of Cot-1 DNA as an RNA FISH probe [29, 30]. Cot-1 DNA (which contains highly repetitive sequences) is routinely used to block non-specific hybridization of genomic probes to repeats, and has been developed as a probe to detect global gene expression using FISH [30]. Cot-1 RNA hybridization provides a convenient assay to identify silent heterochromatic regions within nuclei by the absence of a hybridization signal [30]. Our model for ASAR function predicts that ASAR lncRNAs expressed from every chromosome are detected by Cot-1 RNA FISH due to the presence of repetitive sequences, including abundant L1 antisense sequences, within the ASAR transcripts [26]. This interpretation is consistent with the observation that Cot-1 RNA is comprised predominantly of L1 sequences and is associated with euchromatin throughout interphase nuclei [31]. Furthermore, L1 RNA is localized to interphase chromosome territories, is excluded from heterochromatin, and is associated with the euchromatin fraction of chromosomes even following prolonged transcriptional inhibition [31]. We previously found that ectopic integration of an *ASAR6* transgene leads to loss of Cot1 RNA on the integrated chromosome, suggesting that the ASAR6 transgene silenced the endogenous ASARs on the integrated chromosome [26]. Therefore, because *ASAR6* and *ASAR6-141* are expressed from opposite homologs, our model includes reciprocal silencing of each other in *cis,* resulting in the reciprocal pattern of monoallelic expression of *ASAR6* and *ASAR6-141* on the two chromosome 6 homologs (Fig. 6).

One hallmark of genes that are subject to PRME is coordination in the asynchronous replication between alleles [7–9, 20]. This coordination can be either in *cis*, i.e. the early replicating alleles of two genes are always on the same homolog; or in *trans*, i.e. the early replicating alleles are always on opposite homologs [20]. In this report, we found that the asynchronous replication of *ASAR6-141* is coordinated in *cis* with *ASAR6*. This observation is consistent with our previous findings that human chromosome 6 contains loci that display random asynchronous replication that is coordinated both in *cis* and in *trans*, that some of these asynchronous loci are separated by >100 megabases of genomic DNA, and that the coordinated loci are on either side of the centromere [6, 20]. It will be interesting to determine if all human autosome pairs display a similar coordination in expression and asynchronous replication of PRME genes.

Asynchronous replication of random monoallelic genes is an epigenetic mark that appears before transcription and is thought to underlie the differential expression of the two alleles of identical sequence [18]. Therefore, because the asynchronous replication at PRME genes is coordinated along each chromosome, the expression pattern of PRME genes is also anticipated to be coordinated, i.e. in *cis*- always expressed from the same homolog; or in *trans*- always expressed from opposite homologs. We previously found that *ASAR6* and *ASAR15* are expressed from the later replicating alleles [7, 20]. In contrast, the *FUT9* protein coding gene, which is closely linked to *ASAR6* (see Fig. S2), is expressed from the early replicating allele [7, 20]. Therefore, PRME genes can be expressed from either the early or the late replicating alleles. One unanticipated result from our allelic expression and asynchronous replication assays described here is that *ASAR6-141* is expressed from the early replicating allele, which is the first example of an ASAR that is expressed from the early replicating allele. Nevertheless, we found that disruption of the expressed allele, but not the silent allele of *ASAR6-141* results in delayed replication of chromosome 6, indicating that expression and not asynchronous replication is a critical component of ASAR function. This conclusion is consistent with our previous observation that ASAR6 RNA mediates the chromosome-wide effects of *ASAR6* forced expression [26]. Therefore, the role of asynchronous replication at ASAR loci may serve as a mechanism to help establish which allele will be transcribed. Thus, the epigenetic mark that establishes early and late replication between the two alleles of PRME genes may function to establish asymmetry between alleles, and then depending on the promoter/enhancer elements at different PRME genes either the early or late replicating allele will be transcribed.

One striking feature of both *ASAR6* and *ASAR15* is that they contain a high density L1 retrotransposons, constituting ∼40% and ∼55% of the expressed sequence, respectively [6, 7]. L1s were first implicated in monoallelic expression when Dr. Mary Lyon proposed that L1s represent “booster elements” that function during the spreading of X chromosome inactivation [32, 33]. In humans, the X chromosome contains ∼27% L1 derived sequence while autosomes contain ∼13% [34]. In addition, L1s are present at a lower concentration in regions of the X chromosome that escape inactivation, supporting the hypothesis that L1s serve as signals to propagate inactivation along the X chromosome [34]. Further support for a role of L1s in monoallelic expression came from the observation that L1s are present at a relatively high local concentration near both imprinted and random monoallelic genes located on autosomes [35]. L1s have also been linked to DNA replication timing from the observation that differentiation-induced replication timing changes are restricted to AT rich isochores containing high L1 density [36]. Another potential link between L1s and DNA replication is the observation that ∼25% of origins in the human genome were mapped to L1 sequences [37]. While this observation is suggestive of a relationship between origins and L1s, it is not clear what distinguishes L1s with origin activity from L1s without [37].

During our genetic characterization of *ASAR6* we mapped an ∼29 kb critical region that when deleted results in DRT/DMC [6, 20]. This ∼29 kb region contains one full length and 5 truncated L1s. Similarly, we mapped an ∼124 kb critical region within *ASAR15* that contains 3 full length and 15 truncated L1s [7]. We recently used ectopic integration of transgenes and CRISPR/Cas9-mediated chromosome engineering and found that L1 sequences, oriented in the antisense direction, mediate the chromosome-wide effects of *ASAR6* and *ASAR15* [38]. In addition, we found that oligonucleotides targeting the antisense strand of the one full length L1 within ASAR6 RNA restored normal replication timing to mouse chromosomes expressing an *ASAR6* transgene. These results provided the first direct evidence that L1 antisense RNA plays a functional role in replication timing of mammalian chromosomes [38].

We previously proposed a model in which the antisense L1 sequences function to suppress splicing, and to promote stable association of the RNA with the chromosome territories where they are transcribed [26]. Consistent with this interpretation is the finding that a de novo L1 insertion, in the antisense orientation, into an exon of the mouse *Nr2e3* gene results in inefficient splicing, accumulation of the transcript to high levels, and retention of the transcript at the mutant *Nr2e3* locus [39]. In addition, a more recent study found that the antisense strand of L1 RNA functions as a multivalent “hub” for binding to numerous nuclear matrix and RNA processing proteins, and that the L1 antisense RNA binding proteins repress splicing and 3’ end processing within and around the L1s [40].

The vlincRNAs were identified as RNA transcripts of >50 kb of contiguous RNA-seq reads that have no overlap with annotated protein coding genes [23]. The vlincRNAs were identified from the FANTOM5 Cap Analysis of Gene Expression (CAGE) dataset, indicating that the vlincRNAs contain 5’ caps and consequently represent RNA Pol II transcripts [23]. We previously found that *ASAR6* and *ASAR15* are also transcribed by RNA Pol II [6, 7]. ASAR6-141 RNA shares certain characteristics with ASAR6 and ASAR15 RNAs that distinguish them from other canonical RNA Pol II lncRNAs. Thus, even though ASAR6-141, ASAR6 and ASAR15 RNAs are RNA Pol II products they show little or no evidence of splicing or polyadenylation and remain associated with the chromosome territories where they were transcribed ([6, 7, 23]; and see Fig. 2 and S2). Our work supports a model where all mammalian chromosomes express “ASAR” genes that encode chromosome associated lncRNAs that control the replication timing program in *cis*. In this model, the ASAR lncRNAs function to promote proper chromosome replication timing by controlling the timing of origin firing. In addition, because both *ASAR6,* and *ASAR6-141* are monoallelically expressed, our model includes expression of different ASAR genes from opposite homologs ([26]; see Fig. 6).

We previously found that 5% of chromosome translocations, induced by two different mechanisms (IR and Cre/loxP) display DRT/DMC [4, 5]. Because ∼5% of translocations display DRT/DMC and only one of the two translocation products has DRT/DMC [5], these results indicate that ∼2.5% of translocation products display DRT/DMC [4–7]. Taken with the observation that the translocations that display DRT/DMC have disrupted ASAR genes [6, 7], suggests that ∼2.5% of the genome is occupied by ASARs. The vlincRNAs were identified as nuclear, non-spliced, non-polyadenylated transcripts of >50 kb of contiguously expressed sequence that are not associated with protein coding genes [23]. There are currently >2,700 annotated human vlincRNAs, and they are expressed in a highly cell type-specific manner [23–25]. Because many of the vlincRNAs are encoded by regions of the genome that do not overlap with protein coding genes, many of the vlincRNAs contain a high density of repetitive elements, including L1s (see Fig. 1 and Table S1 for examples). In this report, we found that the genomic region annotated as vlinc273 has all of the physical and functional characteristics that are shared with *ASAR6* and *ASAR15*, and therefore vlinc273 is an ASAR (designated here as *ASAR6-141*). In addition, while ASAR6 RNA was not annotated as a vlincRNA in any previous publication, our RNA-seq data from HTD114 cells indicates that ASAR6 RNA has all of the characteristics of a vlincRNA (see Fig. S2). Furthermore, we note that there are two annotated vlincRNAs (vlinc253 and vlinc254) that map within the ∼1.2 mb domain of *cis*-coordinated asynchronous replication that we previously associated with the *ASAR6* locus ([20]; Fig. S2). Therefore, vlinc253 and vlinc254 display asynchronous replication that is coordinated with *ASAR6*, *ASAR6-141* and all other PRME genes on human chromosome 6 (see [20]). Taken together, these observations raise the intriguing possibility that these other vlincRNAs are also ASARs. Finally, the clustering of vlincRNA genes with ASAR characteristics, and their apparent tissue-restricted expression patterns (see Fig. 1 and S2), supports a model in which each autosome contains clustered ASAR genes, and that these ASAR clusters, expressing different ASAR transcripts in different tissues, function as “Inactivation/Stability Centers” that control replication timing, monoallelic gene expression, and structural stability of each chromosome.

## Methods

### Cell culture

HTD114 cells are a human *APRT* deficient cell line derived from HT1080 cells [41], and were grown in DMEM (Gibco) supplemented with 10% fetal bovine serum (Hyclone). GM12878 cells were obtained from ATCC and were grown in RPMI 1640 (Life Technologies) supplemented with 15% fetal bovine serum (Hyclone). Primary blood lymphocytes were isolated after venipuncture into a Vacutainer CPT (Becton Dickinson, Franklin Lakes, NJ) per the manufacturer’s recommendations and grown in 5 mL RPMI 1640 (Life Technologies) supplemented with 10% fetal bovine serum (Hyclone) and 1% phytohemagglutinin (Life Technologies). All cells were grown in a humidified incubator at 37°C in a 5% carbon dioxide atmosphere.

### RNA-seq

Nuclei were isolated from HTD114 cells following lysis in 0.5% NP40, 140 mM NaCl, 10 mM Tris-HCl (pH 7.4), and 1.5 mM MgCl_2_. Nuclear RNA was isolated using Trizol reagent using the manufacturer’s instructions, followed by DNase treatment to remove possible genomic DNA contamination. RNA-seq was carried out at Novogene. Briefly, ribosomal RNAs were removed using the Ribo-Zero kit (Illumina), RNA was fragmented into 250-300bp fragments, and cDNA libraries were prepared using the Directional RNA Library Prep Kit (NEB). Paired end sequencing was done on a NoaSeq 6000. Triplicate samples were merged and aligned to the human genome (hg19) using the STAR aligner [42] with default settings. Duplicate reads and reads with map quality below 30 were removed with SAMtools [43].

### DNA FISH

Mitotic chromosome spreads were prepared as described previously [2]. After RNase (100µg/ml) treatment for 1h at 37^°^C, slides were washed in 2XSSC and dehydrated in an ethanol series and allowed to air dry. Chromosomal DNA on the slides was denatured at 75^°^C for 3 minutes in 70% formamide/2XSSC, followed by dehydration in an ice cold ethanol series and allowed to air dry. BAC and Fosmid DNAs were labeled using nick translation (Vysis, Abbott Laboratories) with Spectrum Orange-dUTP, Spectrum Aqua-dUTP or Spectrum Green-dUTP (Vysis). Final probe concentrations varied from 40-60 ng/µl. Centromeric probe cocktails (Vysis) and/or whole chromosome paint probes (Metasystems) plus BAC or Fosmid DNAs were denatured at 75^°^C for 10 minutes and prehybridized at 37^°^C for 10 minutes. Probes were applied to denatured slides and incubated overnight at 37^°^C. Post-hybridization washes consisted of one 3-minute wash in 50% formamide/2XSSC at 40^°^C followed by one 2-minute rinse in PN (0.1M Na_2_HPO_4_, pH 8.0/2.5% Nonidet NP-40) buffer at RT. Coverslips were mounted with Prolong Gold antifade plus DAPI (Invitrogen) and viewed under UV fluorescence (Olympus).

### ReTiSH

We used the ReTiSH assay essentially as described [19]. Briefly, unsynchronized, exponentially growing cells were treated with 30μM BrdU (Sigma) for 6 or 5 and 14 hours. Colcemid (Sigma) was added to a final concentration of 0.1 μg/mL for 1 h at 37°C. Cells were trypsinized, pelleted by centrifugation at 1,000 rpm, and resuspended in prewarmed hypotonic KCl solution (0.075 M) for 40 min at 37°C. Cells were pelleted by centrifugation and fixed with methanol-glacial acetic acid (3:1). Fixed cells were drop gently onto wet, cold slides and allowed to air-dry. Slides were treated with 100μg/ml RNAse A at 37°C for 10 min. Slides were rinsed briefly in H_2_0 followed by fixation in 4% formaldehyde at room temperature for 10 minutes. Slides were incubated with pepsin (1 mg/mL in 2N HCl) for 10 min at 37°C, and then rinsed again with H_2_0 and stained with 0.5 μg/μL Hoechst 33258 (Sigma) for 15 minutes. Slides were flooded with 200μl 2xSSC, coversliped and exposed to 365-nm UV light for 30 min using a UV Stratalinker 2400 transilluminator (Stratagene). Slides were rinsed with H_2_0 and drained. Slides were incubated with 100μl of 3U/μl of ExoIII (Fermentas) in ExoIII buffer for 15 min at 37°C. The slides were then processed directly for DNA FISH as described above, except with the absence of a denaturation step. *ASAR6* DNA was detected with BAC RP11-767E7, and *ASAR6-141* DNA was detected with BAC RP11-715D3.

### RNA-DNA FISH

Cells were plated on glass microscope slides at ∼50% confluence and incubated for 4 hours in complete media in a 37°C humidified CO_2_ incubator. Slides were rinsed 1X with sterile RNase free PBS. Cell Extraction was carried out using ice cold solutions as follows: Slides were incubated for 30 seconds in CSK buffer (100mM NaCl/300mM sucrose/3mM MgCl_2_/10mM PIPES, pH 6.8), 10 minutes in CSK buffer/0.1% Triton X-100, followed by 30 seconds in CSK buffer. Cells were then fixed in 4% paraformaldehyde in PBS for 10 minutes and stored in 70% EtOH at −20°C until use. Just prior to RNA FISH, slides were dehydrated through an EtOH series and allowed to air dry. Denatured probes were prehybridized at 37°C for 10 min, applied to non-denatured slides and hybridized at 37°C for 14-16 hours. Post-hybridization washes consisted of one 3-minute wash in 50% formamide/2XSSC at 40^°^C followed by one 2-minute rinse in 2XSSC/0.1% TX-100 for 1 minute at RT. Slides were then fixed in 4% paraformaldehyde in PBS for 5 minutes at RT, and briefly rinsed in 2XSSC/0.1% TX-100 at RT. Coverslips were mounted with Prolong Gold antifade plus DAPI (Invitrogen) and slides were viewed under UV fluorescence (Olympus). Z-stack images were generated using a Cytovision workstation. After capturing RNA FISH signals, the coverslips were removed, the slides were dehydrated in an ethanol series, and then processed for DNA FISH, beginning with the RNase treatment step, as described above.

### Replication timing assay

The BrdU replication timing assay was performed as described previously on exponentially dividing cultures and asynchronously growing cells [44]. Mitotic chromosome spreads were prepared and DNA FISH was performed as described above. The incorporated BrdU was then detected using a FITC-labeled anti-BrdU antibody (Roche). Coverslips were mounted with Prolong Gold antifade plus DAPI (Invitrogen), and viewed under UV fluorescence. All images were captured with an Olympus BX Fluorescent Microscope using a 100X objective, automatic filter-wheel and Cytovision workstation. Individual chromosomes were identified with either chromosome-specific paints, centromeric probes, BACs or by inverted DAPI staining. Utilizing the Cytovision workstation, each chromosome was isolated from the metaphase spread and a line drawn along the middle of the entire length of the chromosome. The Cytovision software was used to calculate the pixel area and intensity along each chromosome for each fluorochrome occupied by the DAPI and BrdU (FITC) signals. The total amount of fluorescent signal in each chromosome was calculated by multiplying the average pixel intensity by the area occupied by those pixels. The BrdU incorporation into human chromosome 6 homologs containing CRISPR/Cas9 modifications was calculated by dividing the total incorporation into the chromosome with the smaller chromosome 6 centromere (6B) divided by the BrdU incorporation into the chromosome 6 with the larger centromere (6A) within the same cell. Boxplots were generated from data collected from 8-12 cells per clone or treatment group. Differences in measurements were tested across categorical groupings by using the Kruskal-Wallis test [45] and listed as P-values for the corresponding plots.

### CRISPR/Cas9 engineering

Using Lipofectamine 2000, according to the manufacturer’s recommendations, we co-transfected HTD114 cells with plasmids encoding GFP, sgRNAs and Cas9 endonuclease (Origene). Each plasmid encoded sgRNAs were designed to bind at the indicated locations (Fig. 1; also see Table S1). 48h after transfection, cells were plated at clonal density and allowed to expand for 2-3 weeks. The presence of deletions in were confirmed by PCR using the primers described in Supplemental Table S1. The single cell colonies that grew were analyzed for heterozygous deletions by PCR. We used retention of a heterozygous SNPs (see Table S1) to identify the disrupted allele (CHR6A vs CHR6;B), and homozygosity at this SNP confirmed that cell clones were homogenous.

## Supporting Information

**S1 Fig.** Delayed replication of chromosome 6 following disruption of *ASAR6*. (A and B) A representative mitotic spread from BrdU (green) treated cells containing a deletion of the expressed allele of *ASAR6* [26]. Mitotic cells were subjected to DNA FISH using a chromosome 6 centromeric probe (red). The larger centromere resides on the chromosome 6 with the expressed *ASAR6* allele (CHR6A). Bar is 10 uM. (C) The two chromosome 6s were extracted from a and b and aligned to show the BrdU incorporation and centromeric signals. (D) Pixel intensity profiles of BrdU incorporation and DAPI staining along the (6A) and (6B) chromosomes from panel C. The long (q) and short (p) arms of chromosome 6 are indicated. Bar is 2 uM. (E) BrdU quantification along 6A and 6B from panel D. (F) The ratio of DNA synthesis into the two chromosome 6s was calculated by dividing the BrdU incorporation in 6B by the incorporation in 6A.

**S2 Fig. Expression and asynchronous replication of *ASAR6*.** UCSC Genome Browser view of the *ASAR6* RNA-seq data, showing the reads from the plus and minus strand in separate tracks, from HTD114 nuclear ribo-minus RNA. We previously mapped the ∼1.2 mb asynchronous replication domain associated with *ASAR6* as indicated (see [20]). We note that the RNA-seq reads associated with *ASAR6* fulfill all of the characteristics described for vlincRNAs (see [23, 24]). Two additional vlincRNAs (253 and 254) also map to the asynchronous replication domain. Also shown is the Repeat Masker Track.

**S1 Table. Repetitive elements within vlinc273 (ASAR6-141).** The chromosome position, orientation, size and total bases occupied by LINE, Alu, and other repeats, from RepeatMasker are shown.

**S2 Table. DNA oligonucleotides used for sgRNAs and PCR primers.** The DNA sequence of the oligonucleotides and the position on chromosome 6 of the oligonucleotides used in for sgRNAs and PCR primers used to screen for deletions. Also shown are the heterozygous SNPs within the PCR products used to determine which allele was expressed and/or deleted following CRISPR/Cas9 expression.

## References

1. Thayer MJ. Mammalian chromosomes contain cis-acting elements that control replication timing, mitotic condensation, and stability of entire chromosomes. Bioessays. 2012;34(9):760–70. PubMed PMID: 22706734.

2. Smith L, Plug A, Thayer M. Delayed Replication Timing Leads to Delayed Mitotic Chromosome Condensation and Chromosomal Instability of Chromosome Translocations. Proc Natl Acad Sci U S A. 2001;98:13300–5.

3. Chang BH, Smith L, Huang J, Thayer M. Chromosomes with delayed replication timing lead to checkpoint activation, delayed recruitment of Aurora B and chromosome instability. Oncogene. 2007;26(13):1852–61. PubMed PMID: 17001311.

4. Breger KS, Smith L, Turker MS, Thayer MJ. Ionizing radiation induces frequent translocations with delayed replication and condensation. Cancer Research. 2004;64:8231–8.

5. Breger KS, Smith L, Thayer MJ. Engineering translocations with delayed replication: evidence for cis control of chromosome replication timing. Hum Mol Genet. 2005;14(19):2813–27. PubMed PMID: 16115817.

6. Stoffregen EP, Donley N, Stauffer D, Smith L, Thayer MJ. An autosomal locus that controls chromosome-wide replication timing and mono-allelic expression. Hum Mol Genet. 2011;20:2366–78. PubMed PMID: 21459774.

7. Donley N, Smith L, Thayer MJ. ASAR15, A cis-Acting Locus that Controls Chromosome-Wide Replication Timing and Stability of Human Chromosome 15. PLoS Genet. 2015;11(1):e1004923. Epub 2015/01/09. doi: 10.1371/journal.pgen.1004923. PubMed PMID: 25569254.

8. Ensminger AW, Chess A. Coordinated replication timing of monoallelically expressed genes along human autosomes. Hum Mol Genet. 2004;13(6):651–8. PubMed PMID: 14734625.

9. Singh N, Ebrahimi FA, Gimelbrant AA, Ensminger AW, Tackett MR, Qi P, et al. Coordination of the random asynchronous replication of autosomal loci. Nat Genet. 2003;33(3):339–41. PubMed PMID: 12577058.

10. Gendrel AV, Marion-Poll L, Katoh K, Heard E. Random monoallelic expression of genes on autosomes: Parallels with X-chromosome inactivation. Semin Cell Dev Biol. 2016. Epub 2016/04/23. doi: 10.1016/j.semcdb.2016.04.007. PubMed PMID: 27101886.

11. Gendrel AV, Attia M, Chen CJ, Diabangouaya P, Servant N, Barillot E, et al. Developmental dynamics and disease potential of random monoallelic gene expression. Dev Cell. 2014;28(4):366–80. Epub 2014/03/01. doi: 10.1016/j.devcel.2014.01.016. PubMed PMID: 24576422.

12. Chess A. Mechanisms and consequences of widespread random monoallelic expression. Nat Rev Genet. 2012;13(6):421–8. PubMed PMID: 22585065.

13. Alexander MK, Mlynarczyk-Evans S, Royce-Tolland M, Plocik A, Kalantry S, Magnuson T, et al. Differences between homologous alleles of olfactory receptor genes require the Polycomb Group protein Eed. J Cell Biol. 2007;179(2):269–76. PubMed PMID: 17954609.

14. Li SM, Valo Z, Wang J, Gao H, Bowers CW, Singer-Sam J. Transcriptome-wide survey of mouse CNS-derived cells reveals monoallelic expression within novel gene families. PLoS One. 2012;7(2):e31751. PubMed PMID: 22384067.

15. Lin M, Hrabovsky A, Pedrosa E, Wang T, Zheng D, Lachman HM. Allele-biased expression in differentiating human neurons: implications for neuropsychiatric disorders. PLoS One. 2012;7(8):e44017. PubMed PMID: 22952857.

16. Bartolomei MS. Genomic imprinting: employing and avoiding epigenetic processes. Genes Dev. 2009;23(18):2124–33. PubMed PMID: 19759261.

17. Gimelbrant A, Hutchinson JN, Thompson BR, Chess A. Widespread monoallelic expression on human autosomes. Science. 2007;318(5853):1136–40. PubMed PMID: 18006746.

18. Mostoslavsky R, Singh N, Tenzen T, Goldmit M, Gabay C, Elizur S, et al. Asynchronous replication and allelic exclusion in the immune system. Nature. 2001;414(6860):221–5. PubMed PMID: 11700561.

19. Schlesinger S, Selig S, Bergman Y, Cedar H. Allelic inactivation of rDNA loci. Genes Dev. 2009;23(20):2437–47. PubMed PMID: 19833769.

20. Donley N, Stoffregen EP, Smith L, Montagna C, Thayer MJ. Asynchronous Replication, Mono-Allelic Expression, and Long Range Cis-Effects of ASAR6. PLoS Genet. 2013;9(4):e1003423. PubMed PMID: 23593023.

21. Kapranov P, St Laurent G, Raz T, Ozsolak F, Reynolds CP, Sorensen PH, et al. The majority of total nuclear-encoded non-ribosomal RNA in a human cell is ’dark matter’ un-annotated RNA. BMC biology. 2010;8:149. Epub 2010/12/24. doi: 10.1186/1741-7007-8-149. PubMed PMID: 21176148; PubMed Central PMCID: PMCPMC3022773.

22. St Laurent G, Savva YA, Kapranov P. Dark matter RNA: an intelligent scaffold for the dynamic regulation of the nuclear information landscape. Front Genet. 2012;3:57. Epub 2012/04/28. doi: 10.3389/fgene.2012.00057. PubMed PMID: 22539933; PubMed Central PMCID: PMCPMC3336093.

23. St Laurent G, Vyatkin Y, Antonets D, Ri M, Qi Y, Saik O, et al. Functional annotation of the vlinc class of non-coding RNAs using systems biology approach. Nucleic Acids Res. 2016;44(7):3233–52. Epub 2016/03/24. doi: 10.1093/nar/gkw162. PubMed PMID: 27001520; PubMed Central PMCID: PMCPMC4838384.

24. St Laurent G, Shtokalo D, Dong B, Tackett MR, Fan X, Lazorthes S, et al. VlincRNAs controlled by retroviral elements are a hallmark of pluripotency and cancer. Genome Biol. 2013;14(7):R73. Epub 2013/07/24. doi: 10.1186/gb-2013-14-7-r73. PubMed PMID: 23876380; PubMed Central PMCID: PMCPMC4053963.

25. Caron M, St-Onge P, Drouin S, Richer C, Sontag T, Busche S, et al. Very long intergenic non-coding RNA transcripts and expression profiles are associated to specific childhood acute lymphoblastic leukemia subtypes. PLoS One. 2018;13(11):e0207250. Epub 2018/11/16. doi: 10.1371/journal.pone.0207250. PubMed PMID: 30440012; PubMed Central PMCID: PMCPMC6237371.

26. Platt EJ, Smith L, Thayer MJ. L1 retrotransposon antisense RNA within ASAR lncRNAs controls chromosome-wide replication timing. J Cell Biol. 2017. Epub 2017/12/31. doi: 10.1083/jcb.201707082. PubMed PMID: 29288153.

27. Goldmit M, Bergman Y. Monoallelic gene expression: a repertoire of recurrent themes. Immunol Rev. 2004;200:197–214. PubMed PMID: 15242406.

28. Galupa R, Heard E. X-Chromosome Inactivation: A Crossroads Between Chromosome Architecture and Gene Regulation. Annu Rev Genet. 2018;52:535–66. Epub 2018/09/27. doi: 10.1146/annurev-genet-120116-024611. PubMed PMID: 30256677.

29. Creamer KM, Lawrence JB. XIST RNA: a window into the broader role of RNA in nuclear chromosome architecture. Philosophical transactions of the Royal Society of London Series B, Biological sciences. 2017;372(1733). Epub 2017/09/28. doi: 10.1098/rstb.2016.0360. PubMed PMID: 28947659; PubMed Central PMCID: PMCPMC5627162.

30. Hall LL, Byron M, Sakai K, Carrel L, Willard HF, Lawrence JB. An ectopic human XIST gene can induce chromosome inactivation in postdifferentiation human HT-1080 cells. Proc Natl Acad Sci U S A. 2002;99(13):8677–82. PubMed PMID: 12072569.

31. Hall LL, Carone DM, Gomez AV, Kolpa HJ, Byron M, Mehta N, et al. Stable C0T-1 repeat RNA is abundant and is associated with euchromatic interphase chromosomes. Cell. 2014;156(5):907–19. Epub 2014/03/04. doi: 10.1016/j.cell.2014.01.042. PubMed PMID: 24581492; PubMed Central PMCID: PMCPmc4023122.

32. Lyon MF. X-chromosome inactivation: a repeat hypothesis. Cytogenet Cell Genet. 1998;80(1-4):133–7. PubMed PMID: 9678347.

33. Lyon MF. The Lyon and the LINE hypothesis. Semin Cell Dev Biol. 2003;14(6):313–8. PubMed PMID: 15015738.

34. Bailey JA, Carrel L, Chakravarti A, Eichler EE. Molecular evidence for a relationship between LINE-1 elements and X chromosome inactivation: the Lyon repeat hypothesis. Proc Natl Acad Sci U S A. 2000;97(12):6634–9. Epub 2000/06/07. PubMed PMID: 10841562; PubMed Central PMCID: PMCPMC18684.

35. Allen E, Horvath S, Tong F, Kraft P, Spiteri E, Riggs AD, et al. High concentrations of long interspersed nuclear element sequence distinguish monoallelically expressed genes. Proc Natl Acad Sci U S A. 2003;100(17):9940–5. PubMed PMID: 12909712.

36. Hiratani I, Leskovar A, Gilbert DM. Differentiation-induced replication-timing changes are restricted to AT-rich/long interspersed nuclear element (LINE)-rich isochores. Proc Natl Acad Sci U S A. 2004;101(48):16861–6. Epub 2004/11/24. doi: 10.1073/pnas.0406687101. PubMed PMID: 15557005; PubMed Central PMCID: PMCPMC534734.

37. Bartholdy B, Mukhopadhyay R, Lajugie J, Aladjem MI, Bouhassira EE. Allele-specific analysis of DNA replication origins in mammalian cells. Nat Commun. 2015;6:7051. Epub 2015/05/20. doi: 10.1038/ncomms8051. PubMed PMID: 25987481; PubMed Central PMCID: PMCPMC4479011.

38. Platt EJ, Smith L, Thayer MJ. L1 retrotransposon antisense RNA within ASAR lncRNA genes controls chromosome-wide replication timing. Journal of Cell Biology. 2017;in press. Epub December 29, 2017.

39. Chen J, Rattner A, Nathans J. Effects of L1 retrotransposon insertion on transcript processing, localization and accumulation: lessons from the retinal degeneration 7 mouse and implications for the genomic ecology of L1 elements. Hum Mol Genet. 2006;15(13):2146–56. Epub 2006/05/26. doi: 10.1093/hmg/ddl138. PubMed PMID: 16723373.

40. Attig J, Agostini F, Gooding C, Chakrabarti AM, Singh A, Haberman N, et al. Heteromeric RNP Assembly at LINEs Controls Lineage-Specific RNA Processing. Cell. 2018;174(5):1067–81.e17. Epub 2018/08/07. doi: 10.1016/j.cell.2018.07.001. PubMed PMID: 30078707; PubMed Central PMCID: PMCPMC6108849.

41. Zhu Y, Bye S, Stambrook PJ, Tischfield JA. Single-base deletion induced by benzo[a]pyrene diol epoxide at the adenine phosphoribosyltransferase locus in human fibrosarcoma cell lines. Mutat Res. 1994;321(1-2):73–9.

42. Dobin A, Davis CA, Schlesinger F, Drenkow J, Zaleski C, Jha S, et al. STAR: ultrafast universal RNA-seq aligner. Bioinformatics (Oxford, England). 2013;29(1):15–21. Epub 2012/10/30. doi: 10.1093/bioinformatics/bts635. PubMed PMID: 23104886; PubMed Central PMCID: PMCPMC3530905.

43. Li H, Handsaker B, Wysoker A, Fennell T, Ruan J, Homer N, et al. The Sequence Alignment/Map format and SAMtools. Bioinformatics (Oxford, England). 2009;25(16):2078–9. Epub 2009/06/10. doi: 10.1093/bioinformatics/btp352. PubMed PMID: 19505943; PubMed Central PMCID: PMCPMC2723002.

44. Smith L, Thayer M. Chromosome replicating timing combined with fluorescent in situ hybridization. J Vis Exp. 2012;(70):e4400. Epub 2012/12/29. doi: 10.3791/4400. PubMed PMID: 23271586; PubMed Central PMCID: PMCPMC3567166.

45. Kruskal JB. Multidimensional scaling by optimizing goodness of fit to a nonmetric hypothesis. Psychometrika. 1964;29:1–27.

